# Uncovering an unconventional JAK1/2-STAT3 branch in macrophage IFNγ signaling

**DOI:** 10.1101/2025.08.13.667888

**Authors:** Mees Botman, Josje M.A. Huisman, Alexandra Drakaki, Sanne L. Maas, Agnieszka Witalisz-Siepracka, Dagmar Stoiber, Emiel P.C. van der Vorst, Ricky Siebeler, Marten A. Hoeksema

## Abstract

Interferon-γ (IFNγ) is a key cytokine in immune activation, especially anti-viral responses and driver of macrophage activation. It classically signals via JAK1/2-mediated STAT1 homodimers. Here, we identify an alternative, non-canonical signaling component in which IFNγ simultaneously also activates STAT3. Our results show that IFNγ activates STAT3 rapidly and directly through JAK1 and JAK2. We provide the first evidence that STAT3 can form heterodimers with STAT1 in this context and demonstrate that STAT3 is co-recruited to a subset of IFNγ-induced, STAT1-bound regulatory elements. While IFNγ directly activates STAT3, our results reveal that its contribution to gene regulation is limited, indicating that STAT1 easily substitutes the STAT1-STAT3 heterodimer for STAT1 homodimers when STAT3 is absent. These findings uncover STAT3 as a new unconventional player in macrophage IFNγ signaling, underscoring the complex and context-dependent nature of cytokine signaling networks.

## Introduction

Interferon γ (IFNγ) is a key cytokine in innate and adaptive immunity, particularly in macrophage activation, anti-viral responses and antigen presentation (1). Upon binding its receptor, it is known to signal through Janus kinase (JAK) 1 and JAK2, which subsequently phosphorylate and activate signal transducer and activator of transcription 1 (STAT1). Upon dimerization, STAT1 homodimers translocate to the nucleus, binding the gamma activated sequence (GAS) element, leading to interferon stimulated gene (ISG) expression (2, 3).

The importance of IFNγ signaling is underscored by its conservation across species and roles in a wide variety of human diseases. Although beneficial during viral infections and boosting anti-cancer immunity, this pathway is known to be hyperactive in autoimmune diseases, like lupus and rheumatoid arthritis (4). Therapeutic strategies aimed at modulating cytokine responses in disease often focus on the JAK-STAT signaling pathway. JAK inhibitors have shown high potential in the treatment of (auto-)inflammatory diseases (5), including rheumatoid arthritis (6) and inflammatory bowel disease (7). Initially, pan-JAK inhibitors such as tofacitinib were developed to inhibit JAK activity broadly. More recently, drug development has shifted toward more selective JAK inhibitors that target individual JAK family members to reduce off-target effects (5). This suggests that subtle differences in downstream signaling pathways following activation upon a single cytokine play important roles in fine-tuning specific outcomes and warrants investigations that fully elucidate the potential signaling pathways and outcomes.

Indeed, despite extensive characterization of the JAK-STAT signaling axis in immune cell responses, growing evidence indicates that additional, non-canonical pathways can be activated. One example is the ability of IFNγ to induce formation of the interferon-stimulated gene factor 3 (ISGF3) complex - a heterotrimer composed of STAT1, STAT2, and interferon regulatory factor 9 (IRF9) in macrophages - which is traditionally associated with IFNα, IFNβ and IFNλ signaling instead (8, 9). Understanding these alternative pathways holds significant potential for a better and more specific modulation of immune responses in a (disease-)context-dependent manner.

Extending to these prior observations that IFNγ signaling is not restricted to STAT1 alone, we here demonstrate that IFNγ also activates STAT3 in a direct manner. Through protein, kinase activity, RNA- and ChIP-seq analyses, we uncover a previously unrecognized IFNγ-induced non-canonical STAT3 activation in macrophages. Our results show that IFNγ activates STAT3 rapidly and directly through JAK1 and JAK2. We uncover a previously unrecognized aspect of IFNγ signaling: STAT1 and STAT3 form heterodimers and co-occupy genomic loci, yet STAT3 appears to have a restricted role in IFNγ-dependent gene activation.

## Results

### IFNγ activates STAT3 in a rapid and direct manner

To study overlap and distinctions between JAK-STAT signaling pathways, we stimulated mouse bone marrow-derived macrophages with IFNβ, IFNγ and IL-6 and determined total protein levels and phosphorylation status of STAT1 and STAT3 after 2 hours of incubation with these cytokines. We observed no differences in total STAT1 and STAT3 protein levels and found IFNβ and IFNγ incubation led to phosphorylation of STAT1 (Fig. 1A, Fig. S1A), in line with previous findings (2, 3). IL-6 stimulation instead led to STAT3 phosphorylation, which also fits the current dogma (3). However, IFNβ and IFNγ stimulation also induced STAT3 phosphorylation (Fig. 1A). While STAT3 phosphorylation upon IFNβ stimulation has been reported previously (10, 11), its induction by IFNγ in macrophages has not.

**Figure 1.**
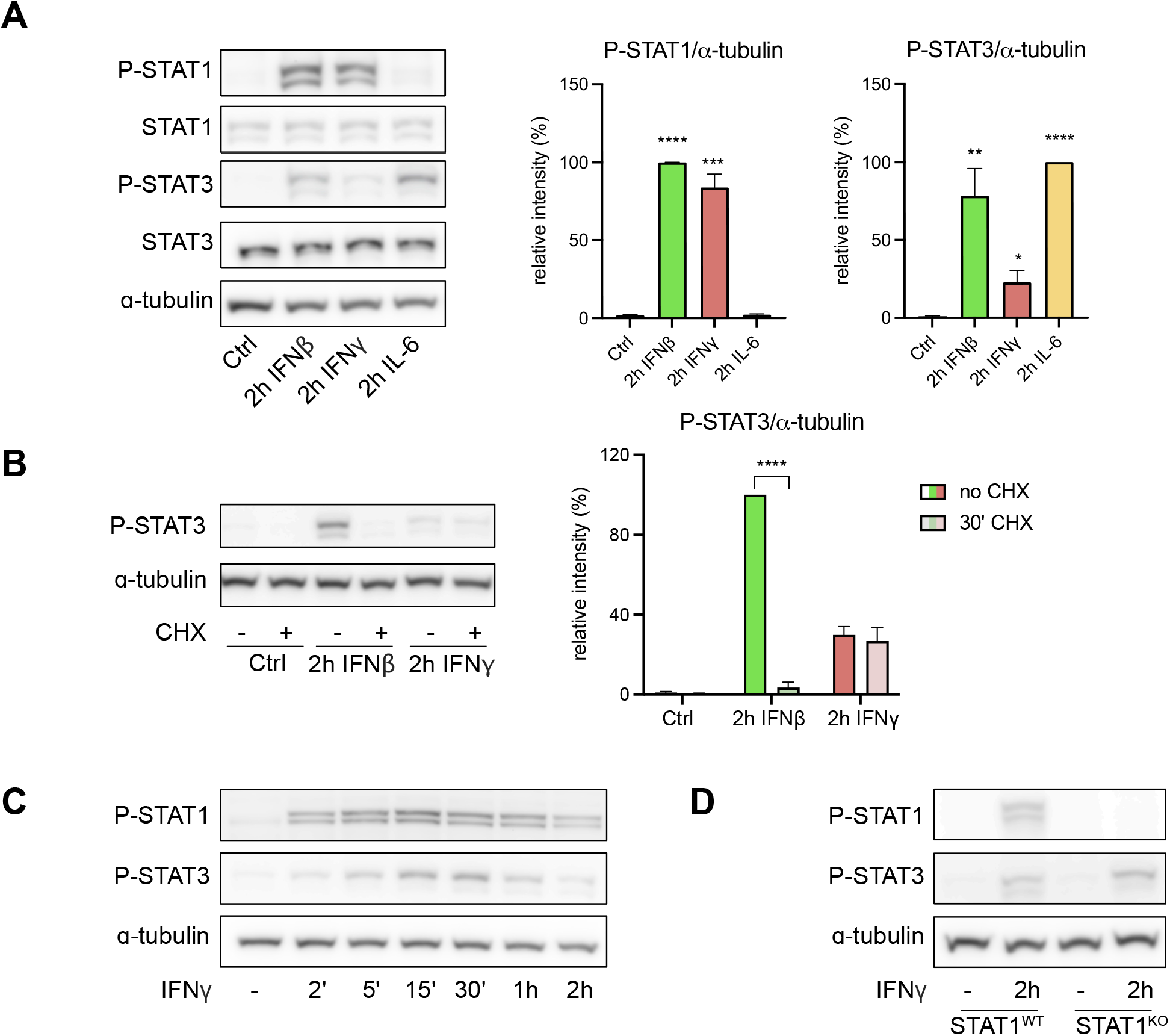
STAT3 activation by IFNs. (A) Mouse bone marrow macrophages, after differentiation for seven days with 15% L929-cell conditioned medium, were stimulated with IFNβ, IFNγ or IL-6 for two hours. (P-)STAT1, (P-)STAT3 and α-tubulin levels were detected using Western Blot (*n*=3). Bars show mean ± SD. (B) Cells were pre-treated with CHX for 30 minutes, after which the macrophages were stimulated with IFNβ or IFNγ for an additional two hours. P-STAT3 and α-tubulin levels were detected using Western Blot (*n*=3). Bars show mean ± SD. Blots were quantified using ImageJ, bars show mean ± SD, *n*=3. One-way ANOVA, * *p* < 0.05, ** *p* < 0.01, \*\*\*\**p* < 0.0001. (C) Time-course experiment in which cells were stimulated for at least two minutes up to two hours with IFNγ (one representative out of three experiments shown). (D) P-STAT1, P-STAT3 and α-tubulin levels were detected in STAT1^WT^ and STAT1^KO^ macrophages stimulated with IFNγ for two hours (one representative out of three experiments shown).

To investigate whether STAT3 phosphorylation takes place directly or indirectly, for example, through the secretion of STAT3-activating cytokines like IL-6 or IL-10 (12, 13), we pre-treated macrophages with cycloheximide (CHX) to block protein translation and, hence, cytokine secretion. Our results indicate that IFNβ-induced STAT3 activation was found to be dependent on protein translation, as CHX pre-treatment significantly decreased IFNβ-induced phosphorylation of STAT3. In contrast, IFNγ-activated STAT3 was not altered by pre-treatment with CHX (Fig. 1B). To further investigate whether STAT3 activation by IFNγ is direct, we performed a time-course experiment with short IFNγ stimulations. We observed that STAT3 is rapidly phosphorylated, starting from two minutes of IFNγ stimulation on, the same as STAT1 phosphorylation (Fig. 1C). Moreover, by using STAT1 knock-out macrophages, we could confirm that STAT3 activation by IFNγ is independent of STAT1 (Fig. 1D), supporting the notion that STAT3 phosphorylation is a direct result of IFNγ receptor signaling. Of note these findings are not specific to mice, as also human monocyte-derived macrophages display STAT3 activation upon IFNγ stimulation (Fig. S1B). Together, these data indicate that IFNγ activates, in addition to STAT1, STAT3, and that this occurs in a rapid and direct manner, independent of *de novo* protein synthesis.

### STAT3 activation through JAK1 and JAK2

To test whether STAT3 activation follows the same signaling route as STAT1 activation upon IFNγ stimulation or through different JAK-(in)dependent routes, we first measured kinase activity upon IFNγ stimulation using a kinase activity assay (Fig. S2A-D, Supplemental Table S1). We found JAK2 and JAK3 to be activated upon 30 minutes of IFNγ stimulation (Fig. 2A). We next tested whether STAT3 activation was indeed JAK-dependent. We pre-treated macrophages with pan-JAK inhibitor tofacitinib for 30 minutes before 2h IFNγ stimulation and confirmed that STAT3 phosphorylation depends on JAK activity (Fig. 2B, Fig. S2E-F). STAT1 activation by IFNγ is thought to take place as a result of JAK1 and JAK2 activity in macrophages (2) and we next questioned whether STAT3 activity is triggered through a different JAK than JAK1 and JAK2, especially since we also measured JAK3 activity in the kinase activity array we performed (Fig. 2A). To test this, we generated knock-outs for all JAKs (JAK1, JAK2, JAK3 and TYK2) in RAW264.7 macrophages. Using CRISPR Cas9 and three different guide RNAs for each JAK, we observed efficient deletion of JAK1-3 and TYK2 (Fig. 2C-F). We revealed that IFNγ-induced STAT3 phosphorylation was independent of JAK3 and TYK2 but depends on JAK1 and JAK2 (Fig. 2C-F) instead. Also, STAT1 activity depends on JAK1 and JAK2, and is independent of JAK3 and TYK2 (Fig. 2C-F). These data indicate that STAT3 activation follows the same signaling pathway as STAT1, while its transcriptional role in IFNγ-stimulated macrophages remains unclear.

**Figure 2.**
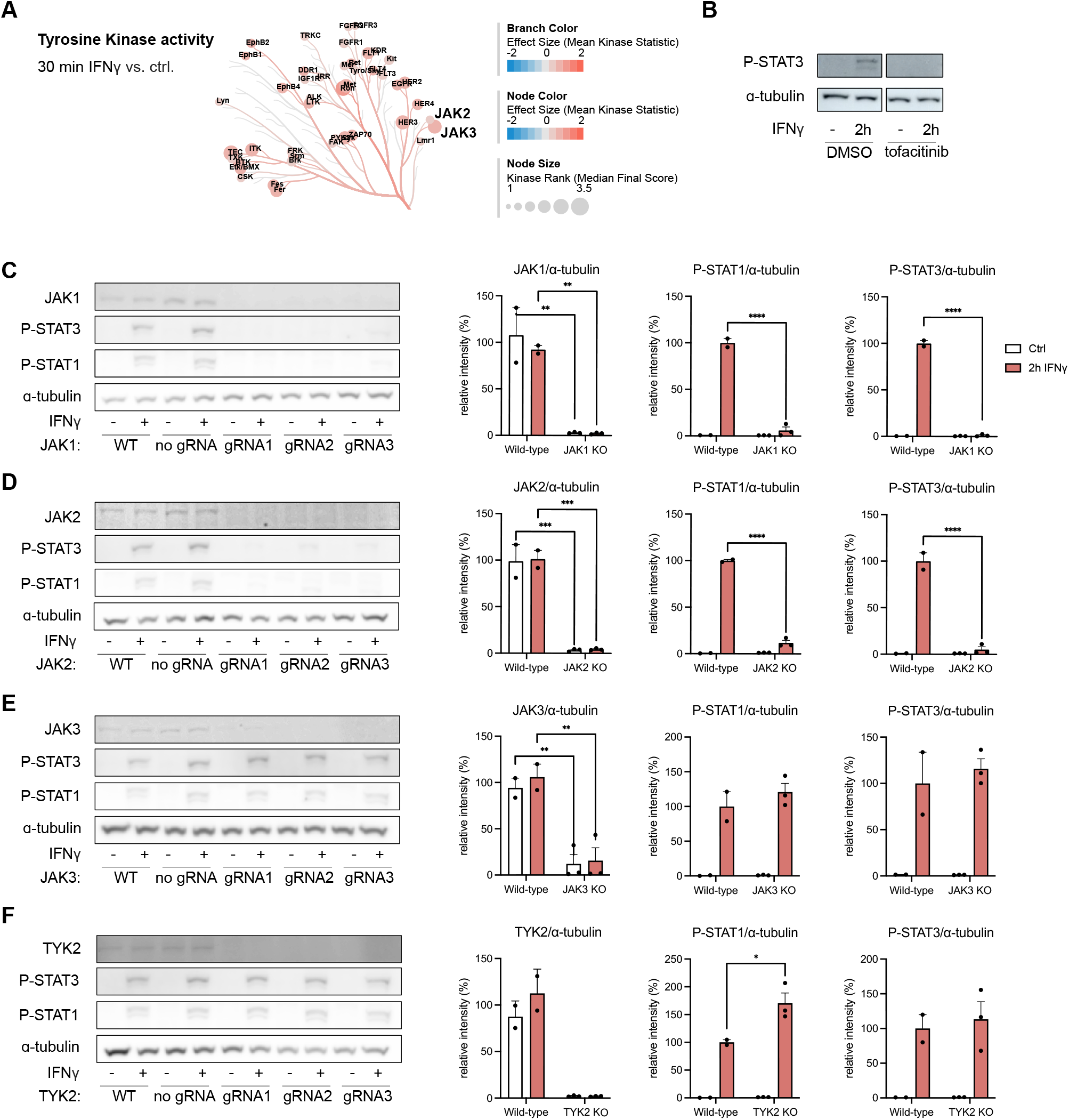
STAT3 activation by IFNγ is JAK1- and JAK2-dependent. (A) Coral activation tree of tyrosine kinase activity array, performed on mouse bone-marrow macrophages that were stimulated with IFNγ for 30 minutes (*n*=4). Full coral activation tree for all tyrosine and serine kinases detected in Figure S2. (B) Cells were pre-treated with tofacitinib for 30 minutes, after which the macrophages were stimulated with IFNγ for an additional two hours. P-STAT3 and α-tubulin levels were detected using Western Blot (one representative out of three experiments shown). (C-F) JAK1-3 and TYK2 knockdown was performed using CRISPR with three different gRNAs per JAK in mouse RAW264.7 cells, no gRNA or WT cells were used as controls. Cells were left unstimulated or stimulated with IFNγ for two hours. JAK1 (C), JAK2 (D), JAK3 (E) and TYK2 (F) were detected using Western Blot as was P-STAT1, P-STAT3 and α-tubulin. Each dot represents a different CRISPR gRNA for KO samples for wild-type it represents untransfected Cas9^+^ RAW264.7 cells or control cells transfected with the same construct, but without gRNAs. Blots were quantified using ImageJ, bars show mean ± SD. Two-way ANOVA, * *p* < 0.05, ** *p* < 0.01, \*\*\*\**p* < 0.0001.

### IFNγ-induced gene expression depends on STAT1, not STAT3

To elucidate whether IFNγ also activates STAT3 target genes, we compared gene expression patterns between IFNγ- and IL-6-stimulated macrophages, as IL-6 is classically known to signal through STAT3 (14). We found a significant overlap in target genes (clusters 5, 6 and 7 in Fig. 3A, Fig. S3A, Supplemental Table S2): 22.8% of the IFNγ-target genes and 84.8% of the IL-6-targets were shared between IFNγ- and IL-6 (Fig. 3B), examples include *Socs3*, a classical IL-6-STAT3 target gene (15). Besides shared target genes, both IFNγ and IL-6 also induced distinctive target gene programs, including *Il10ra* and *Cxcl9* for IFNγ- and *Il10* and *Cd244a* in IL-6-stimulated macrophages (Fig. 3B). Together, this reveals partial convergence between IFNγ and IL-6 signaling pathways.

**Figure 3.**
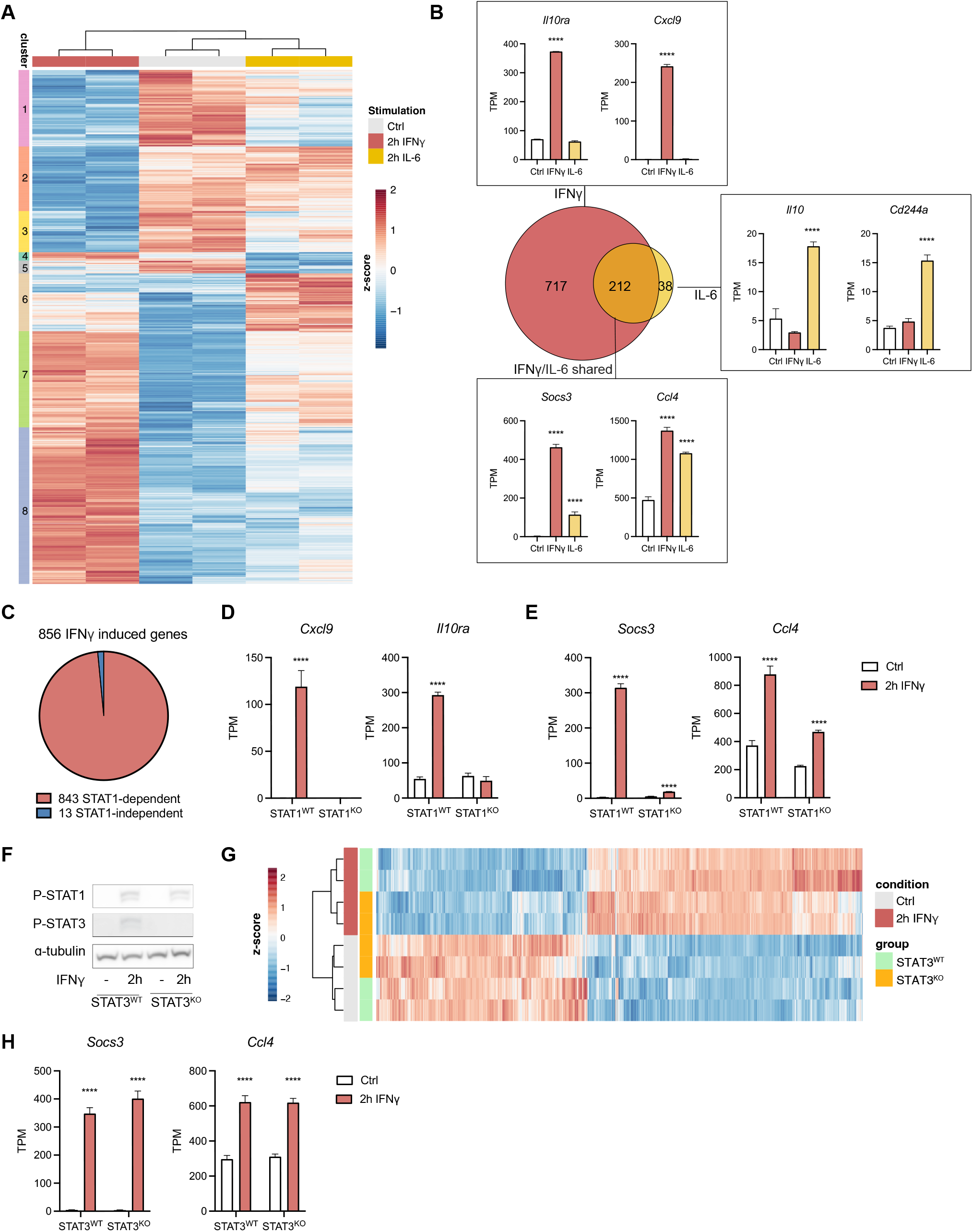
IFN_γ_-induced gene expression depends on STAT1, not STAT3. (A) Heatmap of all differentially expressed genes (DEGs) as measured by RNA-seq, performed on mouse bone marrow macrophages stimulated with IFNγ or IL-6 for two hours, colors represent *z*-score, two replicates per condition. (B) Overlap between IFNγ and IL-6 target genes (logFC > 1, false discovery rate (FDR) < 0.05) with example genes for shared targets and IFNγ- or IL-6-specific target genes. Bars represent mean and SD, TPM = tags per million, **** logFC> 1, FDR < 0.0001). (C) Pie-chart indicating how many IFNγ-upregulated genes in WT (logFC > 1, FDR < 0.05) macrophages are still increased in STAT1^KO^ macrophages. (D) *Cxcl9* and *Il10ra* gene expression in in STAT1^WT^ and STAT1^KO^ macrophages. Bars represent mean and SD, TPM = tags per million, **** logFC > 1, FDR < 0.0001. (E) *Socs3* and *Ccl4* gene expression in in STAT1^WT^ and STAT1^KO^ macrophages. Bars represent mean and SD, TPM = tags per million, **** logFC > 1, FDR < 0.0001. (F) P-STAT1, P-STAT3 and α-tubulin levels were detected in STAT3^WT^ and STAT3^KO^ macrophages stimulated with IFNγ for two hours (one representative out of three experiments shown). (G) Heatmap of all DEGs as measured by RNA-seq, performed on mouse bone marrow STAT3^WT^ and STAT3^KO^ macrophages stimulated with IFNγ for two hours, colors represent *z*-score, two replicates per condition. (H) *Socs3* and *Ccl4* gene expression in in STAT3^WT^ and STAT3^KO^ macrophages, stimulated with IFNγ for two hours, two replicates per condition. Bars represent mean and SD, TPM = tags per million, **** logFC > 1, FDR < 0.0001, compared to control.

To investigate the importance of STAT1 and STAT3 in IFNγ signaling, we first tested whether there is IFNγ-induced gene expression independent from STAT1 activity. Hence, we cultured bone marrow macrophages derived from STAT1^KO^ mice (16). When comparing IFNγ-induced gene expression in STAT1^KO^ macrophages versus wildtype macrophages, we observed that most of the IFNγ-induced gene expression is lost in the absence of STAT1 (Fig. 3C-D, Fig. S3B-C, Supplemental Table S3). However, not all IFNγ target genes are lost, indicating that a subset of genes is activated by IFNγ in a (or at least in part a) STAT1-independent manner (Fig. S3D-E). Amongst others, *Socs3* and *Ccl4* gene expression is still induced in STAT1^KO^ macrophages, albeit at lower levels than in wildtype macrophages (Fig. 3E). As these and other IFNγ-induced genes in STAT1^KO^ macrophages are also induced by IL-6 (Fig. 3B), the data suggest that STAT3 may also play a role in activating these genes.

We subsequently investigated this by determining which IFNγ-target genes were dependent on STAT3. To this end, we performed RNA-seq on *Vav-Cre*^*+*^ *Stat3*^*fl/fl*^ (hematopoietic deletion of *Stat3*, STAT3^KO^ vs *Cre*^*-*^ *Stat3*^*fl/fl*^, STAT3^WT^) bone marrow macrophages stimulated with IFNγ. We found that STAT3 deletion did not affect STAT1 phosphorylation (Fig. 3F), and transcriptome analysis revealed that STAT3 deletion had a minimal impact on IFNγ-induced gene expression (Fig. 3G), with no effect on the previously identified potential STAT3 target genes (Fig. S3F, Supplemental Table S4), like *Socs3* and *Ccl4* (Fig. 3H). Altogether, these data suggest that although STAT3 is activated by IFNγ, just as STAT1, its role is inferior to STAT1.

### STAT1 and STAT3 collaborative binding and recruitment to target genes

To understand the role of STAT3 in IFNγ-stimulated macrophages, we investigated whether it translocates to the nucleus and interacts with STAT1. As previous studies in tumor tissue lysates and T helper cells suggested that STAT1 and STAT3 can form heterodimers (17, 18), we therefore investigated whether such heterodimerization also occurs in macrophages upon IFNγ stimulation. To test this, we performed co-immunoprecipitation (co-IP) assays for P-STAT1 in both unstimulated and IFNγ-stimulated macrophages. Following IFNγ stimulation, we observed robust co-precipitation of P-STAT3 with P-STAT1 (Fig. 4A), indicating that STAT1 and STAT3 physically interact upon IFNγ activation and likely form heterodimers in macrophages.

**Figure 4.**
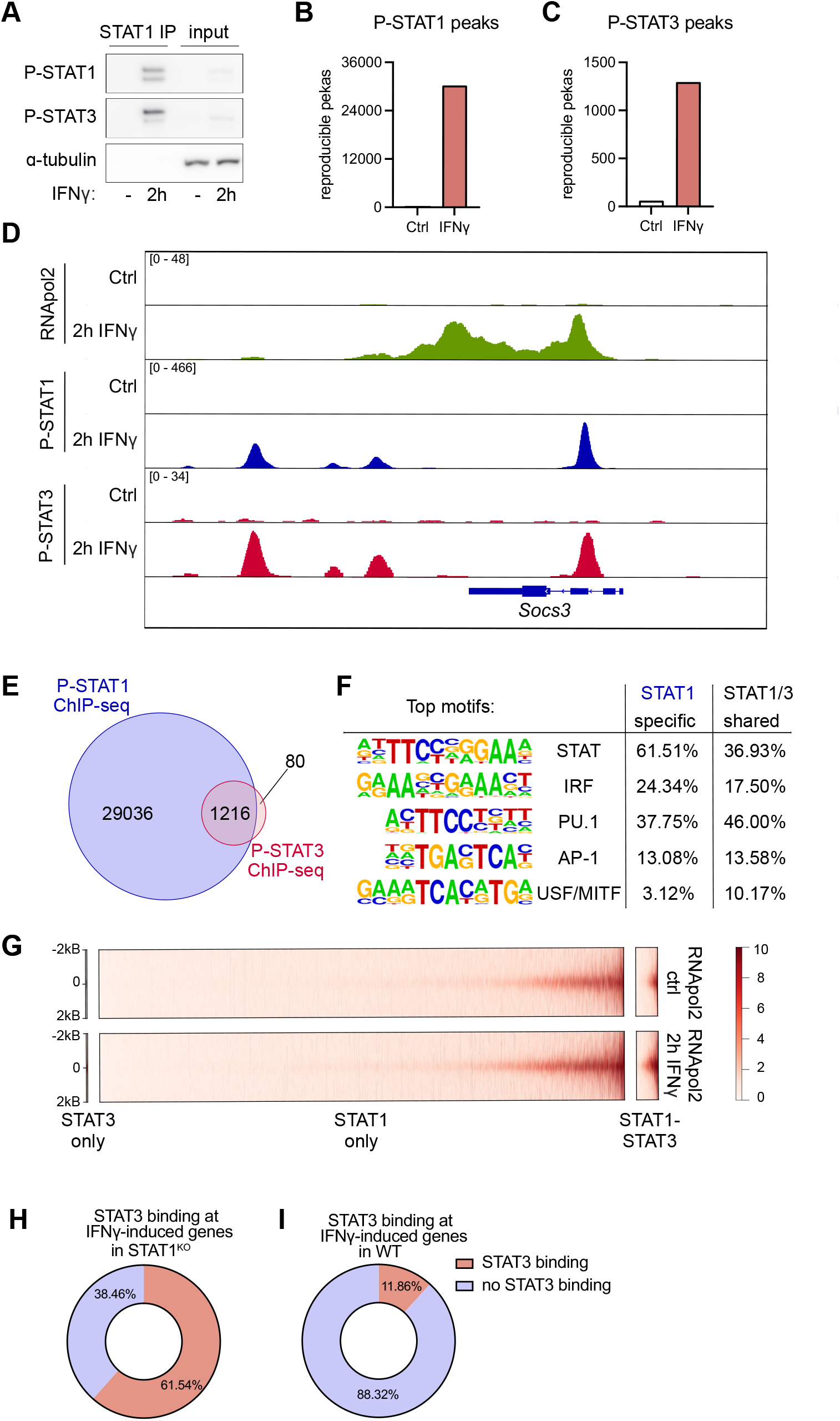
STAT1 and STAT3 collaborative binding and recruitment to target genes. (A) co-IP for P-STAT1 showing association with P-STAT3 as measured with Western Blot, one representative out of three experiments shown, for naïve and IFNγ-stimulated mouse bone marrow macrophages. (B) P-STAT1 ChIP-seq peaks in naïve and IFNγ-stimulated macrophages, bar represents the number of reproducible peaks in two independent ChIP-seq experiments. (C) P-STAT3 ChIP-seq peaks in naïve and IFNγ-stimulated macrophages, bar represents the number of reproducible peaks in two independent ChIP-seq experiments. (D) *Socs3* locus showing enrichment of RNApol2, P-STAT1, P-STAT3 ChIP-seq signal IFNγ-stimulated macrophages, displayed in the IGV genome browser. (E) Overlap in P-STAT1 and P-STAT3 reproducible ChIP-seq peaks in IFNγ-stimulated macrophages. (F) HOMER *de novo* motif analysis on STAT1-specific ChIP peaks versus sites that were bound by both STAT1 and STAT3 based on ChIP-seq signal overlap (from E). (G) RNApol2 ChIP-seq signal intensity at STAT1- or STAT3-specific ChIP peaks versus sites that were bound by both STAT1 and STAT3. (H) STAT3 ChIP-seq peaks at genes that are induced by IFNγ in STAT1^KO^ macrophages. (I) STAT3 ChIP-seq peaks at genes that are induced by IFNγ in STAT1^WT^ macrophages.

To investigate the shared and distinct genomic binding patterns of STAT1 and STAT3 upon IFNγ stimulation, we performed ChIP-seq for P-STAT1 and P-STAT3. In IFNγ-stimulated macrophages, we identified 30,252 P-STAT1 peaks (Fig. 4B) and 1,296 P-STAT3 peaks (Fig. 4C), while binding was minimal under unstimulated conditions. Strikingly, the majority of P-STAT3 peaks overlapped with P-STAT1 peaks (Fig. 4D– E), with only a very limited number of STAT3 unique peaks. This suggests that STAT3 is only recruited to a subset of STAT1-bound loci, for example, the *Socs3* locus, which displays co-occupancy by both factors (Fig. 4D).

Motif analysis revealed that the top five transcription factor binding motifs were highly similar between STAT1-only and STAT1/STAT3-shared peaks, with the major fraction being the STAT motif, also known as the GAS element (Fig. 4F). This makes it difficult to distinguish these groups based on sequence preferences alone. To examine whether co-binding is associated with transcriptional activity, we integrated RNA polymerase II (Pol II) ChIP-seq data. We observed increased Pol II occupancy at both STAT1-only and STAT1/STAT3-shared sites (Fig. 4G), indicating transcriptional engagement at these loci. The *Socs3* locus, again, exemplifies this pattern (Fig. 4D).

As we observed transcriptional activation of a small subset of IFNγ-responsive genes in STAT1^KO^ macrophages, including *Socs3* (Fig. 3C), we investigated whether these STAT1-(partly)-independent genes were enriched for STAT3 binding, as exemplified by the *Socs3* locus (Fig. 4D). Indeed, we found that the majority of genes with STAT1-independent or residual expression showed STAT3 binding (Fig. 4H). In contrast, only 11.7% of all IFNγ target genes in wild-type macrophages displayed STAT3 binding (Fig. 4I).

Together, these findings suggest that STAT3 is co-recruited to a subset of IFNγ-induced, STAT1-bound regulatory elements; but does not establish a distinct genomic binding program. In the absence of STAT1, STAT3 may be capable of binding and activating a select number of genes that are partially STAT1-independent. However, since we did not observe STAT3-dependent transcription, it seems that STAT1 readily substitutes for the STAT1-STAT3 heterodimer through the formation of STAT1 homodimers when STAT3 is absent.

## Discussion

Interferon-γ (IFNγ) is a key cytokine responsible for macrophage activation and anti-viral responses. It primarily signals through the canonical JAK1/2–STAT1 signaling pathway (1-3). In this study, we identify a non-canonical signaling component wherein IFNγ directly and rapidly activates STAT3 through JAK1 and JAK2 activity. We demonstrate that STAT1 and STAT3 can form heterodimers and co-bind a subset of IFNγ-induced regulatory elements. However, loss of STAT3 does not significantly impact the IFNγ-induced transcriptional program, suggesting that STAT3 plays a limited role in gene regulation in this context.

Besides IFNγ, IFNβ stimulation can also lead to STAT3 activation, although indirectly. A likely explanation is that type I IFNs induce secondary cytokines such as IL-10, which is a well-known activator of STAT3. Previous studies have shown that type I IFNs can promote IL-10 expression in macrophages (13). IL-10, in turn, can activate STAT3 in an autocrine or paracrine manner. Previous studies reported STAT3 activation by IFNγ in mouse embryonic fibroblasts (MEFs), but only in the absence of STAT1 (19). Conversely, IL-6, classically signaling through STAT3, has been shown to activate STAT1 in the absence of STAT3 in MEFs (20), further supporting the idea that cytokine signaling is more flexible and context-dependent than previously assumed. The major difference between our findings and the observations in MEFs: in macrophages STAT3 was directly activated by IFNγ, independently of STAT1 presence. We found no evidence for increased STAT3 activation in STAT1-deficient macrophages, nor did we observe enhanced STAT1 activation in the absence of STAT3. These results suggest that, opposed to MEFs, STAT1 and STAT3 are activated simultaneously in macrophages, rather than being compensatory. This suggests that macrophages, as specialized immune cells, engage STAT proteins in a more cell-type-specific and non-redundant manner.

Our kinase activity profiling revealed activation of JAK2 and, unexpectedly, of JAK3 in response to IFNγ stimulation, whereas JAK1 activity was not detected. This contrasts with the well-established role of JAK1 as a critical mediator of IFNγ receptor signaling, suggesting that the lack of detected JAK1 activity may be due to technical limitations of the assay. While JAK2 activation aligns with current models of IFNγ signaling, the observed JAK3 activation is intriguing as it is typically associated with signaling in response to interleukins (21, 22) and its involvement in IFNγ responses has not been reported previously. Our CRISPR knock-out experiments demonstrated that deletion of JAK3 did not impact STAT1 or STAT3 activation upon IFNγ stimulation, indicating that JAK3 is dispensable for IFNγ signaling, leaving it unclear why JAK3 would be activated. Instead, both the canonical STAT1 and unconventional STAT3 activation described here depend on JAK1 and JAK2, reinforcing the central role of these kinases in IFNγ signaling. Since STAT3 phosphorylation in response to IFNγ requires these same JAKs, it indicates that STAT3 activation is part of the intrinsic response towards IFNγ in macrophages.

The presence of STAT1-STAT3 heterodimers and their co-binding at genomic target loci suggests they act collaboratively to regulate transcription. However, despite this physical association, the functional contribution of STAT3 appears limited: deletion of STAT3 does not significantly alter the transcriptional response to IFNγ. This raises the question of the biological relevance of STAT3 activation, heterodimerization with STAT1, and chromatin binding in this context. As STAT1^KO^ macrophages do still show induction of some IFNγ target genes, loci that are enriched for STAT3 ChIP-seq peaks, we propose that STAT1-STAT3 heterodimers are functionally redundant with STAT1 homodimers, in such a way that in the absence of STAT3, STAT1 alone can fulfill all transcriptional roles. Alternatively, STAT3 may play more subtle or temporarily distinct roles; for example, after prolonged stimulation, contributing to maintenance of chromatin accessibility or transcriptional repression (23). Indeed, prior studies have shown that in THP-1 macrophages, STAT3 activation can repress STAT1-dependent ISG expression (10). Additionally, the partial activation of specific IFNγ target genes in STAT1-deficient macrophages suggests that STAT3 may compensate at a subset of loci, albeit weakly.

Altogether, our study demonstrates that IFNγ induces STAT3 activation via a direct, non-canonical, JAK1/2-dependent pathway in macrophages. We provide the first evidence that STAT3 can form heterodimers with STAT1 in this context. Their co-binding at regulatory elements supports this finding. However, the transcriptional contribution of STAT3 appears limited, indicating that it may fine-tune gene expression in more subtle ways. Nevertheless, this new knowledge provides an important next step towards fully understanding of the responses of macrophages to cytokines, a necessity to provide clues or targets for fine-tuning the immune system when it is hyperactive in certain disease settings.

## Resource availability

### Lead contact

Requests for further information, resources and reagents should be directed to the lead contact, Marten Hoeksema.

### Materials availability

Constructs or primer sequences are available upon request.

### Data and code availability

All data needed to evaluate the conclusions in the paper are present in the paper and/or the supplemental information. The sequence data reported in this paper will be deposited in the NCBI GEO database. This paper does not report any original code. For any additional information needed to reanalyze the data reported in this paper, please contact the lead author.

## Acknowledgments

We thank Dr. Mathieu and Dr. Gysemans for providing bone marrow of the STAT1^KO^ mice. This study was supported by ZonMW (grant no. 04510012110011), a Kemp stipendium from the Genootschap ter bevordering van de Natuur-, Geneesen Heelkunde and the European Union (ERC, CytoMAC, 101076170) to M.A.H. Views and opinions expressed are however those of the author(s) only and do not necessarily reflect those of the European Union or the European Research Council Executive Agency. Neither the European Union nor the granting authority can be held responsible for them. E.P.C.v.d.V. was supported by the Corona Foundation (S199/10084/2021), and by the Deutsche Forschungsgemeinschaft (DFG) (SFB TRR219 – Project-ID 322900939; subproject M07).

## Author contributions

M.B., J.M.A.H., A.D., S.L.M. and R.S. performed and designed the experiments and analyzed the data. A.W., D.S. and E.P.C.v.d.V. contributed reagents, materials and advice. M.A.H. conceived and designed the experiments, provided supervision and wrote the manuscript.

## STAR⍰Methods

### Key resources table

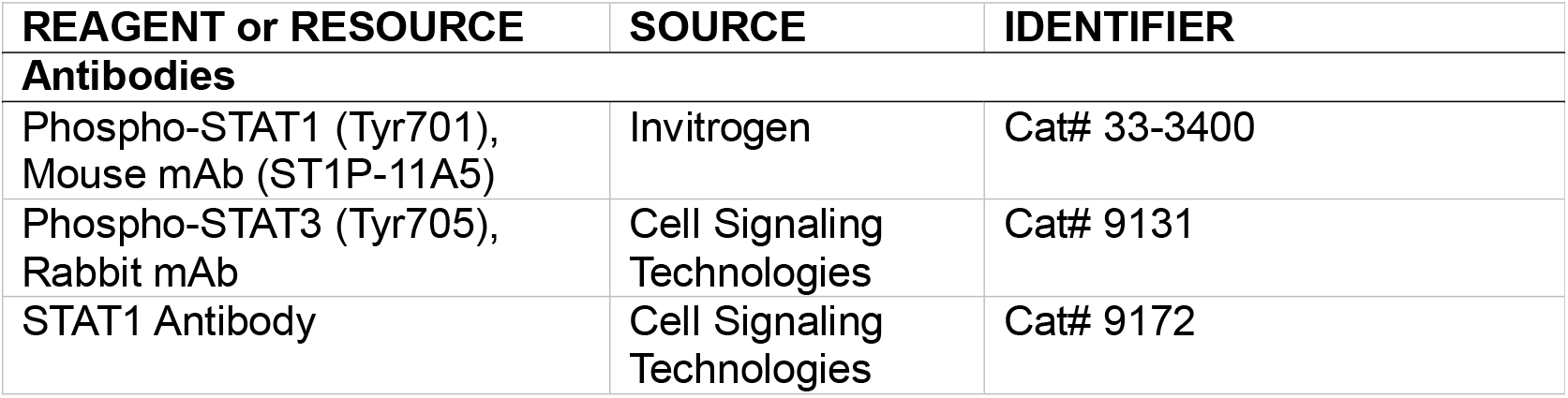
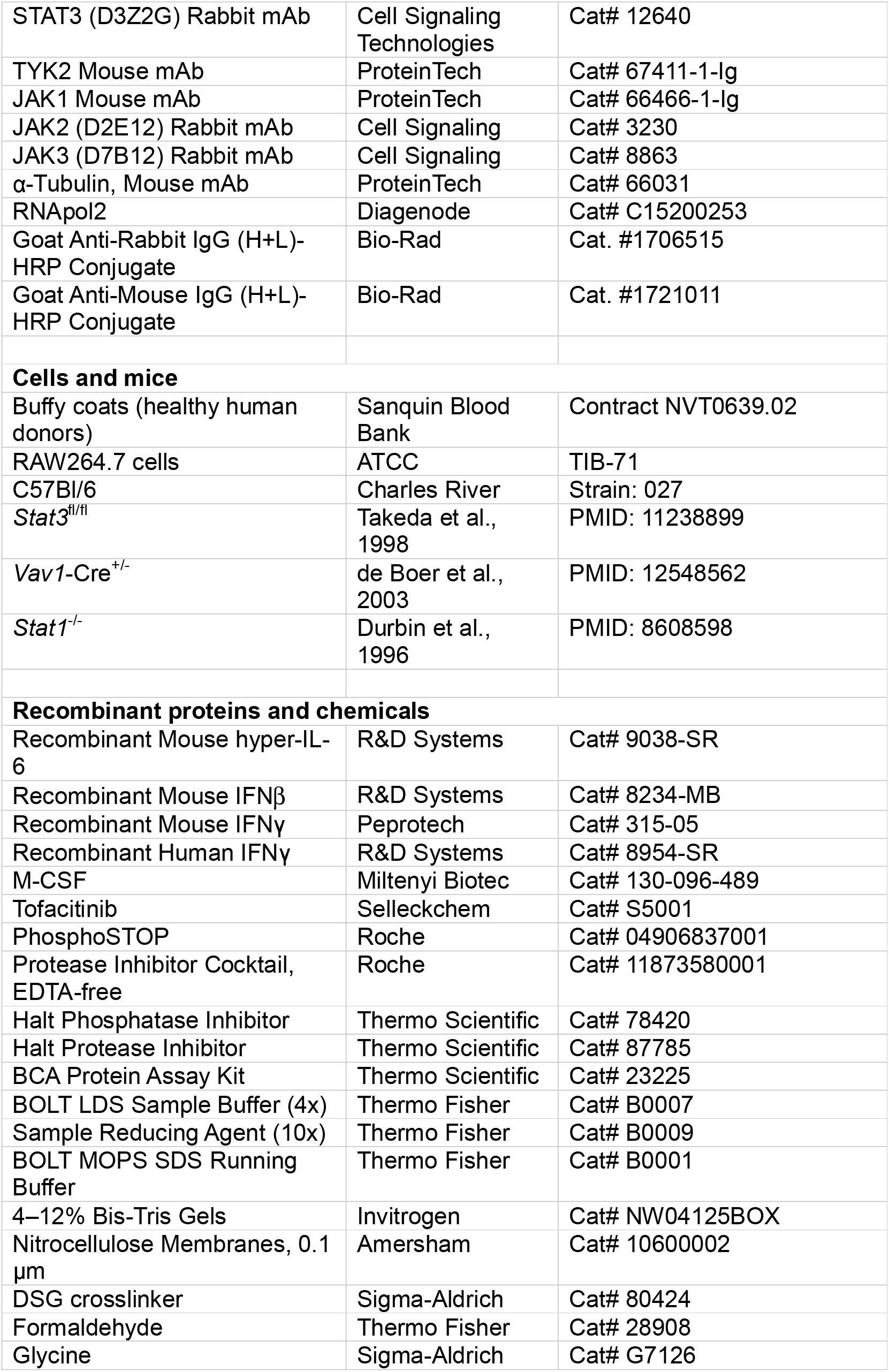
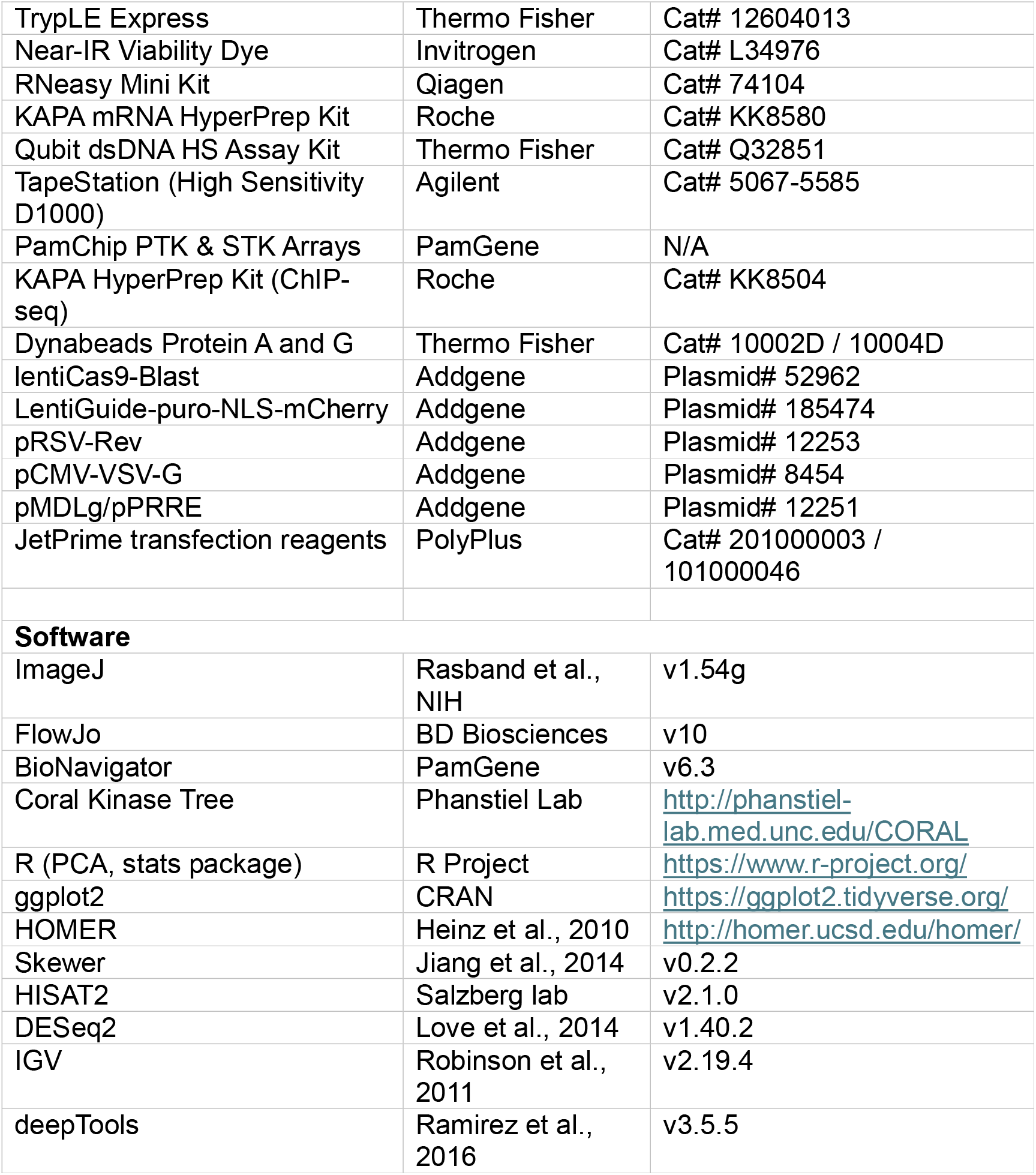

#### Mice

Mice were bred on a C57Bl/6 background and maintained at the Amsterdam UMC animal facility (wild-type mice) and Medical University of Vienna (*Stat3*^*fl/fl*^*VAV1-Cre*^tg/-^ mice) under pathogen-free conditions according to Federation of Laboratory Animal Science Associations guidelines. *Stat3*^*fl/fl*^ mice (24) were crossed with *VAV1-Cre*^*tg*^ mice (25) to obtain *Stat3* deletion in hematopoietic system (*Stat3*^*fl/fl*^*VAV1-Cre*^tg^).

#### Cell lines and cell culture

Bone marrow cells from these wildtype, STAT1^KO^ and STAT3^KO^ mice were differentiated to bone marrow-derived macrophages (BMDMs) using RPMI 1640, 25 mM HEPES, 2 mM L-glutamine, 10% FCS, penicillin (100 U/ml), and streptomycin (100 mg/ml) (all Life Technologies) supplemented with 15% L929-cell conditioned medium (LCM) for 7 days in suspension culture petri dishes. After three days, fresh RPMI medium with 15% LCM was added. Bone marrow macrophages were detached after 7 days of differentiation using TripleE (Life Technologies) and replated in RPMI with 15% LCM for stimulations. Macrophages were stimulated with 50 ng/mL of recombinant mouse IFNβ or IFNγ or 150 ng/mL hyper-IL-6 (R&D systems) for up to two hours. Tofacitinib (pan JAK inhibitor, 50.000 nM, Selleckchem) was added 30 minutes before the IFNγ stimulation. RAW264.7 cells were cultured using DMEM, 25 mM HEPES, 2 mM L-glutamine, 10% FCS, penicillin (100 U/ml), and streptomycin (100 mg/ml) (all Life Technologies) and stimulated with 50 ng/mL of recombinant mouse IFNγ (R&D systems) for two hours.

Peripheral blood mononuclear cells were isolated from healthy donor buffy coats by Ficoll density gradient centrifugation (Lymphoprep, Axishield) and monocytes were isolated using CD14+ bead selection (Miltenyi). Macrophage differentiation from the resulting monocytes was conducted in IMDM (no phenol red) containing 10% FCS, 2 mM L-glutamine, 10% FCS, penicillin (100 U/ml), and streptomycin (all Life Technologies) and 20ng/ml of M-CSF (Miltenyi) for six days and cultured at 37°C and 5% CO2. The whole culture medium was replaced on day three and six. Human macrophages were stimulated with 50 ng/mL of recombinant human IFNγ (R&D systems) for two hours.

#### Western Blot

Cells were lysed in an NP40 lysis buffer (ThermoScientific), containing PhosphoSTOP and protease inhibitor cocktail (Roche). Protein concentrations were measured using BCA (ThermoScientific) and normalized to obtain equal inputs for the Western Blots. Protein samples (10□μg each) were prepared using BOLT SDS/LDS Sample Buffer (4×, Novex) and Sample Reducing Agent (10×, Novex), heated at 70□°C for 10 minutes, and loaded onto pre-cast BOLT Novex 4–12% Bis-Tris gels (10- or 15-well, Invitrogen) with 1× BOLT MOPS SDS Running Buffer. Protein was transferred to Trans-Blot Turbo RTA Mini 0.2 µm Nitrocellulose membranes (Bio-Rad)using the Trans-Blot Turbo system (Bio-Rad) after which the membranes were blocked overnight at 4□°C in TBS with 5% BSA and incubated overnight at 4□°C with primary antibodies: α-tubulin (Proteintech), Phospho-STAT1 (Invitrogen), STAT1 (Cell Signaling), Phospho-STAT3 (Cell Signaling,), STAT3 (Cell Signaling), JAK1 (ProteinTech), JAK2 (Cell Signaling), JAK3 (Cell Signaling) and TYK2 (ProteinTech) all diluted 1:1000 in TBS-1% BSA, except α-tubulin (1:5000). Goat Anti-Mouse and Goat Anti-Rabbit Secondary antibodies (1:5000, Bio-Rad) were applied for 1 hour at room temperature. Detection was performed using chemiluminescence (ImageQuant800, Amersham) and quantified using ImageJ.

#### Kinase Activity Profiling

Cells were lysed on ice for 10 minutes using M-PER Mammalian Protein Extraction Reagent (Thermo Scientific), supplemented with Halt Phosphatase Inhibitor Cocktail and Halt Protease Inhibitor Cocktail, EDTA-free (1:100 each; Thermo Scientific). Lysates were centrifuged at 15,000 × g for 15 minutes at 4°C. Protein concentrations were determined according to the manufacturer’s instructions using the Pierce™ BCA Protein Assay Kit (Thermo Scientific).

Tyrosine kinase (PTK) and serine/threonine kinase (STK) activity profiling was performed using PamChip® peptide kinase microarrays on the PamStation®12 platform (PamGene International). The PTK chip contains 196 phospho-sites, and the STK chip contains 144 phospho-sites.

For PTK profiling, 8.0 μg of protein lysate (n = 4 per condition) was used. The PTK Basic Mix was prepared by combining frozen lysate with 4 μL of 10× PTK reaction buffer, 0.4 μL of 100× BSA, 0.4 μL of 1 M dithiothreitol (DTT), 4 μL of 10× PTK additive, 4 μL of 4 mM ATP, and 0.6 μL of a monoclonal anti-phosphotyrosine FITC-conjugated detection antibody (clone PY20). The final volume was adjusted to 40 μL with distilled water. Before loading the mix onto the chip, a blocking step with 30 μL of 2% BSA was performed, followed by washing with PTK solution. Next, 40 μL of the PTK Basic Mix was added to each chip, and the assay was run for 94 cycles. Fluorescent signal was captured using a CCD camera on the PamStation®12 at kinetic read cycles 32–93 with exposure times of 10, 50, and 200 milliseconds, and at endpoint read cycles using exposures of 10, 20, 50, 100, and 200 milliseconds.

For STK profiling, 1.0 μg of protein lysate (n = 4 per condition) and 400 μM ATP were used per assay. The STK Basic Mix included 1× PTK reaction buffer, 1× BSA, and 1× STK antibody mix. As with PTK assays, a 30 μL blocking step with 2% BSA was performed, followed by washing with PTK solution. The STK Basic Mix (40 μL) was then applied to the array. Samples were incubated at 30°C for 1 hour with dynamic pumping through the porous chip material to enhance binding kinetics and reduce assay time. Phosphorylation was detected using a FITC-conjugated antibody specific for phosphorylated serine/threonine residues.

#### Kinase activity data visualization

Imaging and data analysis were performed using the BioNavigator software version 6.3. Principal Component Analysis (PCA) was conducted to explore patterns of variation and clustering among samples based on peptide phosphorylation profiles. The analysis was performed using the stats package (26) in R, which applies singular value decomposition to the centered and scaled data matrix to examine covariances and correlations between samples. To visualize the results, the gglot2 (27) was used to generate two-dimensional PCA plots, highlighting sample distribution along the principal components. These visualizations enabled the identification of sample grouping, variability, and potential outliers.

To visualize the classification and distribution of kinases identified in the assay, a kinase coral tree was generated. This approach groups kinases based on their evolutionary and functional relationships into families and subfamilies, providing a hierarchical representation of kinase activity profiles. The kinase tree was constructed using the Coral web tool (http://phanstiel-lab.med.unc.edu/CORAL), which allows the mapping of user-defined kinase lists onto a curated phylogenetic tree based on kinase domain homology. Input data included the list of kinases detected as active in the PTK or STK PamChip® assays. The kinases were grouped into their respective families according to the classification system provided by the Human Kinome Project. Visualization parameters were adjusted to reflect relative kinases; here, node/branch colors represent mean kinase statistics (<0.05), and node sizes represent the median final scores (>1.2). The resulting coral tree was used to assess family-wide trends, highlight condition-specific kinase activation patterns, and support the functional interpretation of the dataset.

#### CRISPR

Guide RNAs specific for JAK1 (gRNA 1: TCCGGCTCCACTACCGCATG, gRNA 2: GCTTGGTGCTCTCATCGTAC, gRNA 3: ATGACAACGAACAGTCTGTA), JAK2 (gRNA 1: ATCTTCGCTCGAACGCACTT, gRNA 2: ACGGGACACTCCGTATCTGT, gRNA 3: TCCACATAGACGAGTCAACC), JAK3 (gRNA 1: TGAGCGCGTAGCGGTCGGCC, gRNA 2: GCCTGCGTGTCACGAAGTTC, gRNA 3: CTGGAGACATGTCACCGCTT) and TYK2 (gRNA 1: GGATCTCCTCCTCGCTAGAC, gRNA 2: AAACCACACCGGTATACAGC, gRNA 3: TCTGCGGGACCTGTCTAGCG) were taken from the GeCKOv2 library (28), cloned into the LentiGuide-puro-NLS-mCherry vector (Addgene) and amplified in competent DH5α cells. Extraction of vector-gRNAs was performed with Miniprep (Qiagen) and the ligation was validated by Sanger sequencing. HEK293T cells were transfected with pCMV-VSV-G, pMDLg/PRE, pRSV-Rev (Addgene) and vector-gRNAs or lentiCas9-Blast (Addgene) constructs using JetPrime (Polyplus). Virus was collected and transferred to RAW264.7 cells. After 72 hours, the virus was removed and puromycin (5 μg/ml, Sigma) or blasticidin (10 μg/ml, Sigma) selection was initiated and after colonies observed, cells were further expanded. Virus containing vector-gRNAs were then transferred to Cas9+ RAW264.7 cells. PCRs were performed on isolated genomic DNA (GeneJET, Thermo) to confirm genomic disruption. In short, the DNA regions spanning the sgRNA target site were amplified by PCR. PCR products were analyzed by agarose gel electrophoresis and excised PCR products (Qiagen) were analyzed using Sanger sequencing analysis (AUMC Core Facility Genomics).

#### Co-immunoprecipitation

Cells were lysed on ice using NP40 lysis buffer (ThermoScientific) supplemented with PhosphoSTOP and protease inhibitor cocktail (Roche). Lysates were collected by scraping, clarified by centrifugation (5 □ min, 20,000xg, 4°C) and part of the supernatants were stored as input controls. The remaining lysates were incubated overnight at 4°C with 2 □ μg of P-STAT1 antibody (ThermoScientific) while rotating. On the second day, 20 □ μL of Protein G Dynabeads (after three washes with NP40 lysis buffer containing inhibitors) was added to the antibody-lysate mixtures and incubated for 2 □ hours at room temperature. After this incubation, beads were washed three times and proteins were eluted by incubating with 100 □ μL elution buffer (50 mM Glycine pH=2,75 (Sigma), 1x NuPAGE LDS sample buffer (ThermoScientific) and 1x NuPAGE denaturizing agent (ThermoScientific) at 70 □ °C for 10 □ minutes and magnetic separation. Samples were analyzed using P-STAT1 and P-STAT3 Western Blot as indicated before, alongside input and supernatant fractions to assess pulldown efficiency, to identify protein-protein interactions.

#### RNA isolation and RNA-seq

RNA was isolated of 500,000 cells using the RNeasy Mini kit (Qiagen) according to manufacturer’s instructions, including DNAse treatment, and eluted in 35 μL of elution buffer.

500 ng of total RNA per sample in 25 μL of H2O was used as input for RNA-seq library preparations using half reactions of the KAPA mRNA HyperPrep kit (Roche) according to manufacturer’s instructions. RNA was fragmented at 85C for 6 minutes and UDI barcodes were used for labeling. Library DNA amounts were quantified using Qubit dsDNA HS Assay Kit (Thermo Fisher Scientific), assessed using TapeStation, pooled and 150 bp paired-end sequenced on the NovaSeqXPlus by the AUMC Core Facility Genomics).

#### RNA-seq analysis

Reads were trimmed with Skewer (29)and aligned to mm10 using Hisat2 v2.1.0. Read counts were obtained using featureCounts v2.1.1 (30). TPM values were calculated by normalizing raw read counts to gene length (in kilobases) to obtain Reads Per Kilobase (RPK), scaling RPKs by the sum of all RPKs per sample (in millions), and dividing each RPK by this scaling factor to account for sequencing depth. Differently expressed genes (DEGs) were identified using DESeq2 (31). Significantly regulated genes were defined as expressed genes with p-value adjusted for multiple testing (FDR < 0.05) and log2 fold-change of at least 1.

#### ChIP-seq

ChIP was performed in biological replicates as described previously (32). Briefly, cells were incubated with 2mM DSG (Sigma) in PBS for 30 minutes at RT, followed by 1% formaldehyde (ThermoFischer) for 10 minutes for dual crosslinking, quenched with 0.13M Glycine for 5 minutes and snap-frozen. 6 million cells were sonicated using the BioRuptor Pico sonicator (Diagenode) in 300 μl lysis buffer (10 mM Tris/HCl pH 7.5, 100 mM NaCl, 1 mM EDTA, 0.5 mM EGTA, 0.1% deoxycholate, 0.5% N-laurosylsarcosine, 1× protease inhibitor cocktail). After sonication, Triton X-100 was added to 1% final concentration and lysates were spun at full speed for 10 min; 1% was taken as input DNA, and immunoprecipitation was carried out overnight with 20 μl Protein A Dynabeads (Invitrogen) and 2 μg specific antibodies for RNApol2, P-STAT1 and P-STAT3. Beads were washed three times each with wash buffer I (20 mM Tris/HCl, 150 mM NaCl, 0.1% SDS, 1% Triton X-100, 2 mM EDTA), wash buffer II (10 mM Tris/HCl, 250 mM LiCl, 1% IGEPAL CA-630, 0.7% Na-deoxycholate, 1 mM EDTA), TE 0.2% Triton X-100 and TE 50 mM NaCl and subsequently resuspended 25 μl 10 mM Tris/HCl pH 8.0% and 0.05% Tween-20. ChIP-seq libraries were prepared while bound to Dynabeads using KAPA hyperprep library preparation kit (Roche) using half reactions. DNA was polished, polyA-tailed, and ligated after which dual UDI (IDT) barcodes were ligated to it. Libraries were eluted and crosslinks reversed by adding to the 55 μl KAPA ligation reaction: 22.2 μl water, 5 μl 10% SDS, 5.4 μl 5 M NaCl, 3.75 μl 0.5 M EDTA, 2 μl 0.5 M EGTA, 1.2 μl RNAse (10 mg/ml), and 1.2 μl 20 mg/ml proteinase K, followed by incubation at 55°C for 1 hr and 75°C for 30 min in a thermal cycler. Dynabeads were removed from the library using a magnet and libraries were cleaned up by adding the same volume of Ampure beads as the reaction itself, mixing well, then incubating at room temperature for 10 min. Beads were collected on a magnet and washed two times with 150 μl 80% ethanol for 30s. After the second ethanol wash, beads were air dried and DNA eluted in 15 μl 10 mM Tris/HCl pH 8.0% and 0.05% Tween-20. DNA was amplified by PCR for 14 cycles in a 25 μl reaction volume using KAPA HiFi master mix (Roche) and primer mix. Libraries were size selected using Ampure double-sided bead selection: 0.56-0.80X. Sample concentrations were quantified by Qubit dsDNA HS Assay Kit (Thermo Fisher Scientific) and 150 bp paired-end sequenced on the NovaSeqXPlus by the AUMC Core Facility Genomics.

#### ChIP-seq analysis

Reads were trimmed with Skewer (29) and aligned to mm10 using Hisat2 v2.1.0. BAM files were processed with HOMER’s makeTagDirectory v4.11.1(33) to generate tag directories, which were converted to bigWig coverage files via makeUCSCfile and visualized in IGV (34). Peaks were called from these tag directories using HOMERs findPeaks and reproducible peaks were identified by merging replicate peaks using mergePeaks with default settings. ChIP-seq signal intensity heatmaps and correlation plots were generated using deepTools (35). ChIP-seq signal intensity heatmaps were generated using computeMatrix and visualized with plotHeatmap. To assess global correlations between ChIP-seq datasets, multiBigwigSummary was used in bins mode, followed by plotCorrelation to produce Pearson correlation heatmaps.

#### Cell viability using flow cytometry

After the cells were plated in a 96-well plate and the corresponding treatments were finished, the cells were subjected to flow cytometry to assess cell viability. First, media was removed, and cells were detached using TrypLE Express and neutralized with DMEM. Cells were then spun down at 300xg for 5 minutes at 4 □ °C, supernatant was removed and the cells were washed with PBS. Cells were then stained with the fixable viability dye near-IR (780) fluorescent reactive dye (Invitrogen, 1:1000) diluted in PBS and incubated for 30 minutes at 4 □ °C, protected from light. Finally, cells were washed in FACS buffer (PBS, 0.5% BSA, 0.1% EDTA) and resuspended in 150 □ μL FACS buffer for measurement on the CytoFLEX (Beckman Coulter). Data were analyzed using FlowJo. In FlowJo, viability was determined based on the near-IR viability dye fluorescent signal.

## Figure Legends

**Figure S1, related to Figure 1. STAT3 activation by IFNs**

(A) Mouse bone marrow macrophages, after differentiation for seven days with 15% L929-cell conditioned medium, were stimulated with IFNβ, IFNγ or IL-6 for two hours. STAT1, STAT3 and α-tubulin levels were detected using Western Blot (*n*=3). Bars show mean ± SD.

(B) Human monocyte-derived macrophages, after differentiation for seven days with recombinant M-CSF, were stimulated with IFNγ for two hours. P-STAT1, P-STAT3 and α-tubulin levels were detected using Western Blot (one representative out of three experiments shown).

**Figure S2, related to Figure 2. STAT3 activation by IFN**γ **is JAK1- and JAK2-dependent**.

(A) Coral activation tree of tyrosine kinase activity array, performed on mouse bone-marrow macrophages that were stimulated with IFNγ for 30 minutes (*n*=4).

(B) PCA plot of tyrosine kinase activity array, performed on mouse bone-marrow macrophages that were stimulated with IFNγ for 30 minutes (*n*=4).

(C) Coral activation tree of serine kinase activity array, performed on mouse bone-marrow macrophages that were stimulated with IFNγ for 30 minutes (*n*=4).

(D) PCA plot of tyrosine serine activity array, performed on mouse bone-marrow macrophages that were stimulated with IFNγ for 30 minutes (*n*=4).

(E) Viability staining using flow cytometry on mouse macrophages pre-treated with tofacitinib for 30 minutes, after which the macrophages were stimulated with IFNγ for an additional two hours (*n=6*).

(F) Example of viability staining using flow cytometry on singlet-gated cells.

**Figure S3, related to Figure 3, IFN**_γ_**-induced gene expression depends on STAT1, not STAT3**

(A) PCA plot on the top 500 most variable genes expression data as measured by RNA-seq, performed on mouse bone marrow macrophages stimulated with IFNγ or IL-6 for two hours, two replicates per condition.

(B) PCA plot on the top 500 most variable genes expression data as measured by RNA-seq, performed on STAT1^WT^ and STAT1^KO^ macrophages stimulated with IFNγ for two hours, two replicates per condition.

(C) Heatmap of all differentially expressed genes (DEGs) as measured by RNA-seq, performed on mouse bone marrow STAT1^WT^ and STAT1^KO^ macrophages stimulated with IFNγ for two hours, colors represent *z*-score, two replicates per condition.

(D) Volcano plot on DEGs in STAT1^KO^ macrophages by 2h IFNγ stimulation.

(E) Overlap between IFNγ target genes (logFC > 1, false discovery rate (FDR) < 0.05) in STAT1^WT^ and STAT1^KO^ macrophages stimulated with IFNγ for two hours.

(F) PCA plot on the top 500 most variable genes expression data as measured by RNA-seq, performed on STAT3^WT^ and STAT3^KO^ macrophages stimulated with IFNγ for two hours, two replicates per condition.

**Figure S4, related to Figure 4, STAT1 and STAT3 collaborative binding and recruitment to target genes**

(A) Correlation plots for RNApol2, pSTAT1 and pSTAT3 ChIP-seq.

(B) *Il10ra* locus showing RNApol2, P-STAT1, P-STAT3 ChIP-seq signal in the IGV genome browser.

## Supplemental information

Figures S1-S4.

Supplemental Table S1. Kinase activity array

Supplemental Table S2. TPMs RNA-seq IFNγ vs. IL-6.

Supplemental Table S3. TPMs RNA-seq STAT1^KO^ macrophages.

Supplemental Table S4. TPMs RNA-seq STAT3^KO^ macrophages.

