## Supplementary figures and images for "Uncovering an unconventional JAK1/2-STAT3 branch in macrophage IFNγ signaling"

### Suppl Fig 1

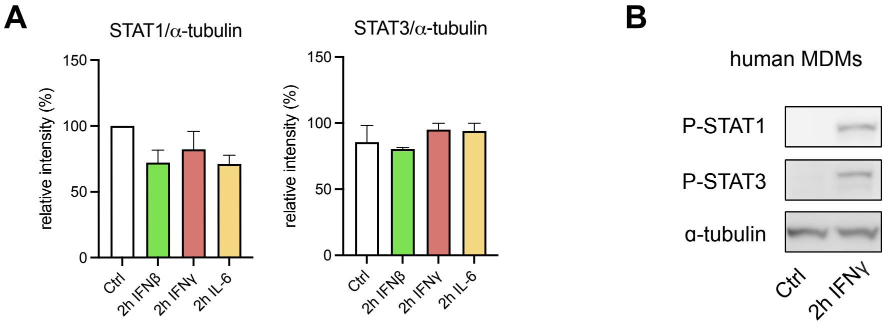

### Suppl Fig 2

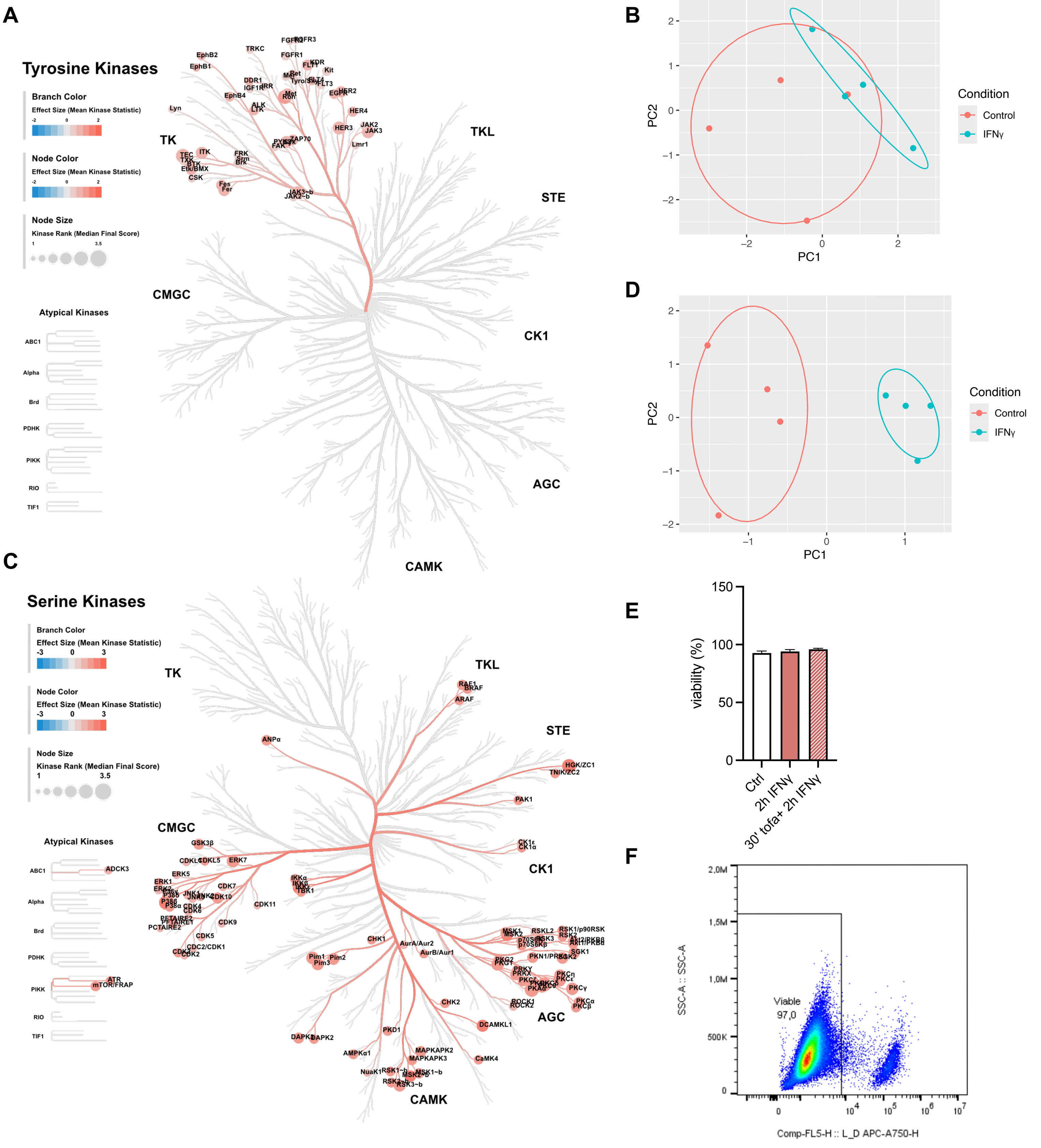

### Suppl Fig 3

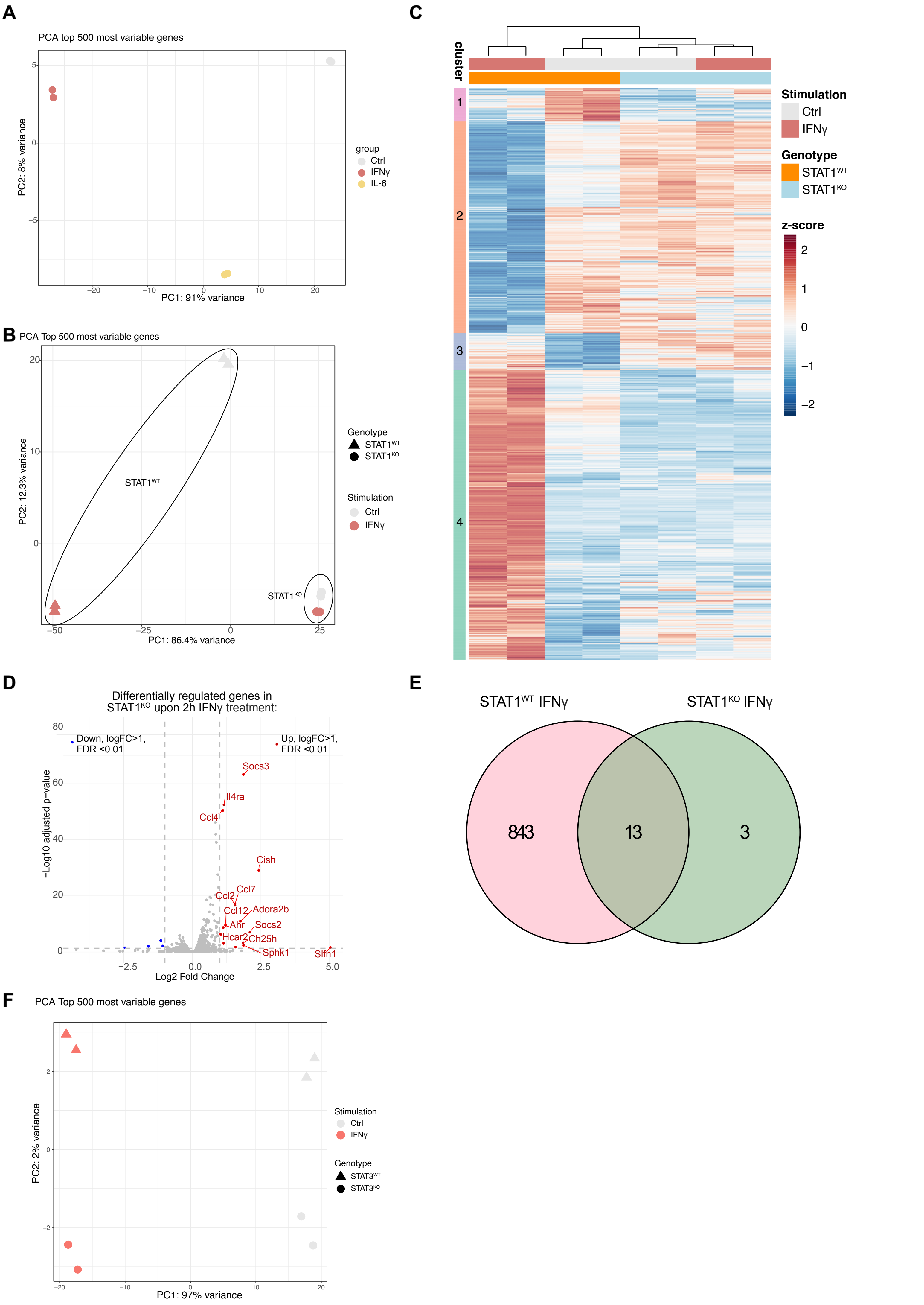

### Suppl Fig 4

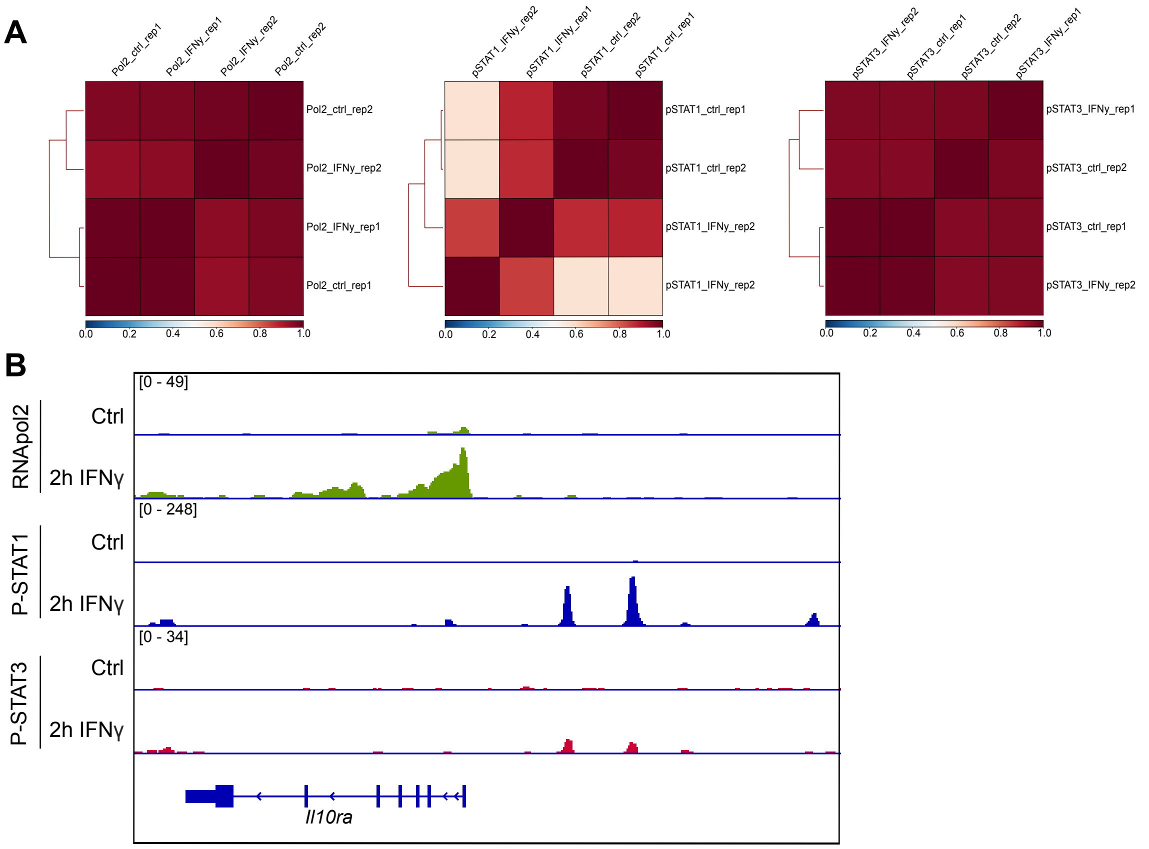
